# Identification and functional categorization of the most stably and most variably expressed genes and retained introns in *Arabidopsis thaliana* seedlings

**DOI:** 10.1101/2024.08.26.608733

**Authors:** Wen-dar Lin, Tatsuo Kanno, Antonius J.M Matzke, Marjori Matzke

## Abstract

We used a large, uniform, RNA-seq dataset to investigate the stability of gene expression and intron retention in *Arabidopsis thaliana* seedlings. Functional classification of the most stably and most variably expressed genes was determined by GO enrichment analyses. Many variably expressed genes encoded proteins involved in photosynthesis and chloroplast structure, likely reflecting their sensitivity to changeable light intensities, and in stress responses, which allow plants to cope with environmental challenges. As revealed by GO enrichment analysis, the most stably expressed genes were involved in protein, lipid and vesicle trafficking, suggesting that fluctuations in expression of these genes are suboptimal for normal seedling development. GO analyses of genes containing either usually retained or usually spliced introns revealed no consistent enrichments for any specific functional categories. However, highly retained introns were often located in the first or last position, which may contain 5’ and 3’ UTRs necessary for transcriptional regulation and mRNA transport and stability. Conversely, usually spliced introns were more frequently located in internal portions of pre-mRNAs, indicating that reliable splicing in coding regions is needed to prevent the formation of premature stop codons. The large RNA-seq dataset we generated can be useful for investigating additional aspects of gene expression and pre-mRNA splicing in Arabidopsis seedlings and sets a precedent for future large scale transcriptome analyses.

## Introduction

The routine use of high throughput nucleic acid sequencing technologies has enabled the generation of a large number of RNA-seq datasets from various plant species, including many from *Arabidopsis thaliana* (Arabidopsis) (Upton *et al*., 2023; Canton *et al*., 2021; Shintani *et al*., 2024). While useful, comparisons of RNA-seq data sets among multiple studies (Zhuo et al., 2016; Yang et al., 2023) are potentially confounded by nonuniformity in plant growth conditions and RNA isolation procedures in individual laboratories.

As the ‘wild-type’ control for transcriptome experiments carried out on various Arabidopsis mutants isolated in our lab (Kanno *et al*., 2020 and references therein), we repeatedly performed RNA-seq on the same control line that was mutagenized to produce these mutants. Control and mutant seedlings for the transcriptome experiments were cultivated under the same sterile growth conditions on solid synthetic medium. Two-weeks after seed sowing, seedlings were harvested and total RNA was prepared at a similar time of day using the same isolation procedure. These efforts resulted in the production of 50 independent RNA-seq datasets that can be directly compared to each other to investigate various aspects of global gene expression in the control line. A previous study in yeast demonstrated that at least six and optimally twelve biological replicates should be used to ensure valid biological interpretation of results from transcriptome experiments (Schurch *et al*., 2016). Our sample size of 50 transcriptomes far exceeds these values and therefore can be expected to provide well-substantiated information.

We used these 50 control RNA-seq datasets to identify and functionally categorize the most stably and most variably (least stably) expressed genes and retained introns in two-week-old Arabidopsis seedlings. GO enrichment analyses of the top 100 genes in each group were performed to assign these genes to specific functional categories. Our aim was to identify the major functional classes that would emerge in the stable and variable groups from our use of a large dataset. Here we report the results of our study and discuss their possible biological implications.

## Methods

### Plant cultivation, RNA isolation procedure and RNA-seq methods

The transgenic plant line used in this study (referred to herein as ‘control’) was created in the *Arabidopsis thaliana* ecotype Col-0 as described in a previous paper (Kanno *et al*. 2008) and recently used in a forward genetic screen using EMS as a mutagen to generate mutants defective in pre-mRNA splicing (Kanno *et al*., 2020).

For seed germination, mutant and control seeds were sterilized by washing in a solution of 0.1 % TritonX/70 % ethanol followed by quickly rinsing in 100 % ethanol and drying on filter paper in a laminar flow hood. Dried seeds were then sprinkled on sterile, solid Murashige and Skoog (MS) medium containing 3 % sucrose in plastic Petri dishes. Following two-days of stratification at 4 °C, seeds were cultivated in a growth incubator under a 16-hour light/8-hour dark cycle at 24°C for two weeks. After this time, total RNA was isolated from about 100 mg of seedlings using a Plant Total RNA Miniprep kit (GMbiolab, Taiwan). Library preparation for RNA-seq was performed as described previously by an in-house Genomic Technology Core facility (Kanno *et al*., 2016).

All RNA-seq FASTQ files were submitted to the NCBI SRA, where the SRA run accessions were numbered from SRR28087940 to SRR28087989. Corresponding BioSamples are under BioProject PRJNA1080259. (reviewer link: https://dataview.ncbi.nlm.nih.gov/object/PRJNA1080259?reviewer=52me2obnjt14d068c901faih8t)

### Using of Ararpot11 and AtRTD3 databases

In this study, we combined Araport11 (Cheng, *et al*., 2017) and AtRTD3 (Zhang, *et al*., 2022) databases as our reference database. AtRTD3 contains 39732 genes and Araport11 38194 genes. AtRTD3 identified 1541 novel gene loci and covered most genes in Araport11, with the exceptions of AT1G48267 and AT5G39693, which were not included in AtRTD3. Additionally, AT2G08855 only appeared in chimeric records. Taking AtRTD3 as the base, and removing chimeric gene records from AtRTD3 and adding the three genes from Araport11, the reference database we used contains non-chimeric records of 39735 genes. Note that AtRTD3 contains no gene category information and we created one genome annotation GFF3 file based on the reference database with gene category information restored from Araport11. The gene categories include mRNA for coding genes and others for non-coding genes. That is, tRNA, rRNA, miRNA_primary_transcript, ncRNA, lncRNA, antisense_lncRNA, snRNA, snoRNA, transcript_region, pseudo_transcript, and pseudo_tRNA were included. uORF and transposable_elements were not included because the former contains no exon intervals and the latter is not closely related with transcription.

### Bioinformatic analysis of RNA-seq transcriptome data

The complete RNA-seq dataset contains approximately 3.5G reads from the 50 samples (see **Figure 1** for size distribution of the dataset). RNA-seq reads were preprocessed using Trimmomatic v0.39 for adapter removal, where reads shorter than 80bps after trimming were dropped. Cleaned reads were mapped to the reference transcriptome database using Bowtie2 v2.2.6 (Langmead and Salzberg, 2012), where only alignments of read pairs mapping to the same transcripts were accepted. Rest reads were mapped to the TAIR10 genome sequences using BLAT (Kent, 2002), where alignment blocks at the two ends were required to be no shorter than 8bps. For the aforementioned two alignment steps, only alignments with identity (number of matches/read length) >=0.95 were accepted. Finally, all reads that could not be mapped to the Arabidopsis genome and transcriptome were mapped to the GFP-containing vector pAJM-EPRV (GenBank accession: HE582394.1) directly using BLAT, only alignments with identity >=0.9 were accepted. In general, more than 95% of reads after trimming were mapped for each sample in the above process. Statistics of the above mapping procedure are available in **Table S1**.

**Figure 1:**
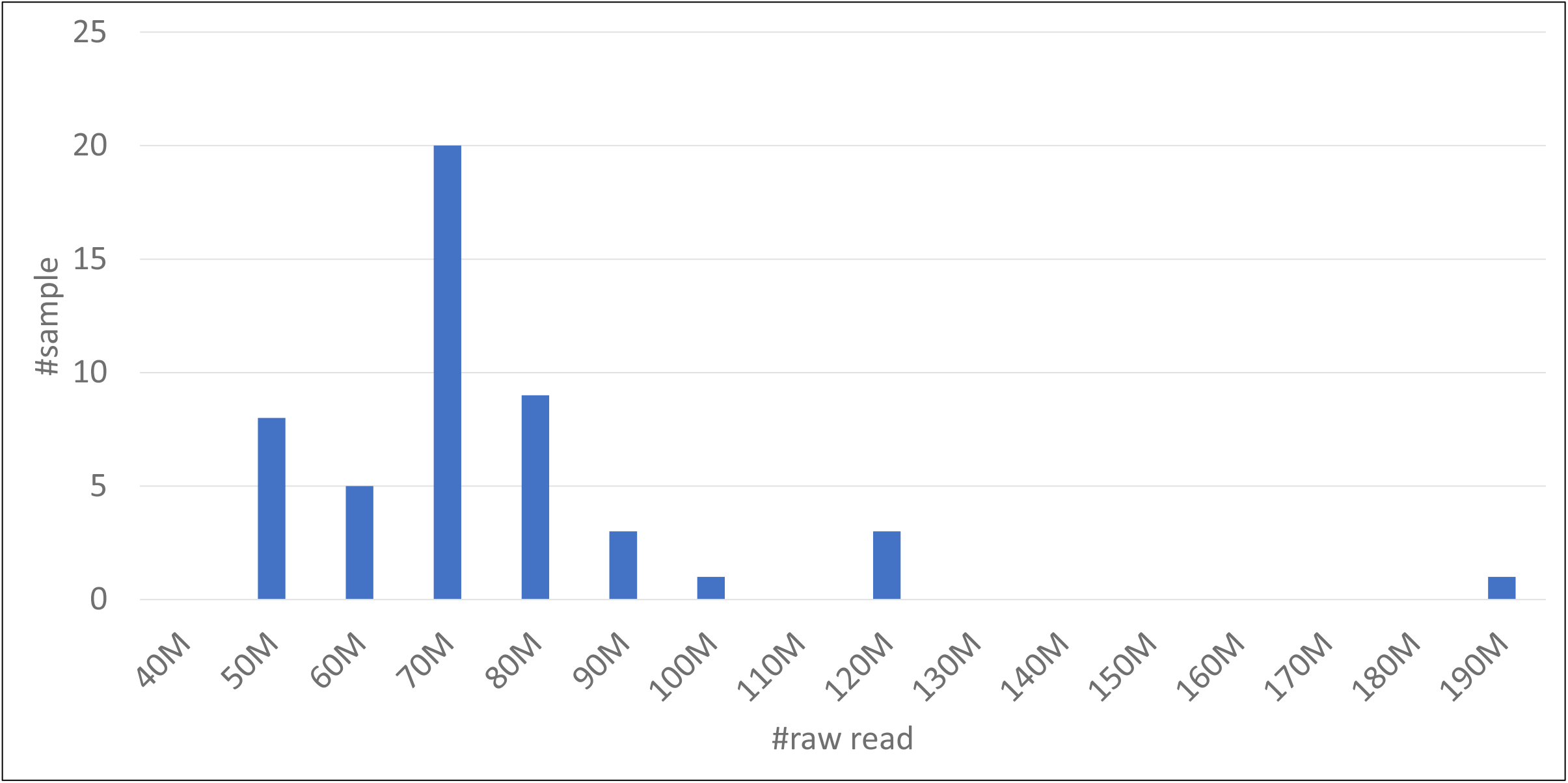
Dataset size distribution of the 50 RNA-seq samples.

#### Gene expression: identification of the least variably and most variably expressed genes

Read count of genes of all samples were computed using the RackJ toolkit (https://sourceforge.net/projects/rackj/), where normalized RPKM (reads per kilo base-pair of gene model per million mapped reads) values were computed based on those read counts normalized by the TMM method (Robinson and Oshlack, 2010). Coefficients of variation (CV, the standard deviation divided by the mean) were computed for all 39735 genes based on the normalized RPKM values of the 50 samples. The 100 genes with smallest CVs were selected as the least variably (most stably) expressed genes. In this group, CVs range from 0.043 to 0.060. (**Table S2**)

For the identification of the most variably expressed genes, 1000 random tests were performed by comparing 25 samples against the other 25 samples using Z-TEST (Kal, *et al*., 1999). For each of the 1000 randomized comparisons, a series of p-value thresholds 0.1, 0.05, 0.01, 0.005, and 0.001 were applied for defining DEGs (differentially expressed genes). Every gene was thus associated with a number of being DEGs under different thresholds. Top 100 genes that were most frequently defined as DEGs were selected as the 100 most variable genes. (**Table S3**)

Note that the application of Z-TEST here is to compare two merged samples; that is, every randomized comparison was aimed to see if the difference between the averages of the two halves was significant or not.

#### Identification of usually spliced (least retained) introns

In the analysis of usually spliced introns, we considered 134815 introns, where each of them was spliced in at least one isoform in our reference database. Intron retention level was determined by comparing read depth of every intron against those of its neighboring exons, where read depths were computed using the RackJ toolkit (https://sourceforge.net/projects/rackj). As read depths of exons were computed based only on uniquely mapped reads, read depths of introns were computed based on both uniquely mapped reads and multiply mapped reads. In so doing, the computation of “usually spliced” would be inferred by considering all reads in the samples but not relying only on unique reads. This should avoid misjudgments on introns being mapped by some multiply mapped reads. Accordingly, for every intron, any of its neighboring exons that showed read depth greater than half depth of the intron in at least 40 samples was identified as a *reference exon*. In so doing, an intron may have none, one, or two reference exons, and we defined its *retention ratio* as read depth of the intron divided by average read depth of its reference exon(s) in a sample. As a result, 4314 introns showed ratios equal to or less than 0.005 in all 50 samples. Further ranking of these *usually spliced* introns was made based on their products of the 50 depth ratios, where zero depths were adjusted to be one tenth of minimum non-zero depths of introns of similar lengths. (**Tables S5 and S6**)

To identify multi-intron genes in which most introns are usually spliced, we collected introns with the aforementioned unadjusted ratios lower than 0.01 in all 50 samples and examined whether any multi-intron genes contain more than 90% of such introns. Thirty genes satisfying the criteria are listed in **Table S7**.

#### Intron retention: identification of the usually retained introns

The identification of usually retained introns is similar to that used for usually spliced introns except that read depth for intron regions and exon regions were both computed based on only uniquely mapped reads using the RackJ toolkit (https://sourceforge.net/projects/rackj/). Accordingly, 471 introns showed retention ratios greater than or equal to 0.9 in all 50 samples. Further ranking of these 471 introns were made based on their products of the 50 retention ratios. These introns were classified as *usually retained* introns. (**Tables S8 and S9**)

Using the 0.9 threshold on retention ratios, we examined whether any multi-intron genes (those with three or more introns) contain more than 90% of introns usually retained. However, no multi-intron genes comprised more than 90% introns showing retention ratios above threshold 0.9, and we lowered the threshold to see if any genes with almost all introns showing retention ratios above some certain threshold. In so doing, we found three multi-intron genes with more than 90% of introns with retention ratios all above 0.7 in all 50 samples. (**Table S10**).

#### Positional trends of usually retained introns and usually spliced introns

To obtain a global view of potential positional preference of usually retained introns and usually spliced introns in multi-intron genes, we made a positional aggregate graph for a given list of introns. To do that, genes were aligned to a 100-slot array by considering their introns evenly separated to the 100 slots. For every intron in the list, we add 1’s to all slots that are corresponding to the intron. For example, the first 25 slots would be added by 1’s if we are counting for the first intron from a four-intron gene. In so doing, a positional trend could be observed if a particular position is generally preferred in multi-intron genes.

#### Gene Ontology enrichment (GO) computation

GO enrichment was done by the following steps to adopt the most updated GO annotation to the date that this study was conducted. Firstly, the basic version of GO database OBO file and the GO annotation GAF file of Arabidopsis thaliana were downloaded from the GO Consortium website (https://geneontology.org/), where their creation dates were marked “2023-01-01”. GO enrichment was done by using GOBU (https://gobu.sourceforge.io/, version 20230301) with an input file created based on the above OBO and GAF files. Enrichment p-values reported in this manuscript were computed using the TopGO elim method (Alexa *et al*., 2006).

## Results

### I. Identification and functional analysis of the least variably and most variably expressed genes in two-week-old Arabidopsis seedlings

#### 100 least variably (most stably) expressed genes (Table S2)

The 100 genes with the lowest coefficients of variation (CV) were identified and analyzed by GO analysis. The most highly ranked terms in the GO_C class (cellular component) were found to be: endosome, trans-Golgi network, clathrin coat of trans-Golgi network vesicle, and Golgi transport complex (ranks 1-4, respectively) (**Figure 2** and **Table S2**). Genes in these categories are involved in intracellular protein and lipid transport, sorting and targeting of proteins by the Golgi complex, as well as vesicle trafficking in the endocytic/endosomal and secretory pathways (Otegui*eet al*.(2010). These functional classes also appeared prominently among the most stably expressed genes in both the GO_C class [(early endosome (rank 10), endoplasmic reticulum (rank 13) and late endosome (rank 21)], and the GO_P (biological process) class, [intracellular protein transport (rank 1), protein targeting to vacuole (rank 2), vacuole organization (rank 7) and Golgi vesicle transport (rank 10)] (**Figure 2** and **Table S2**).

**Figure 2:**
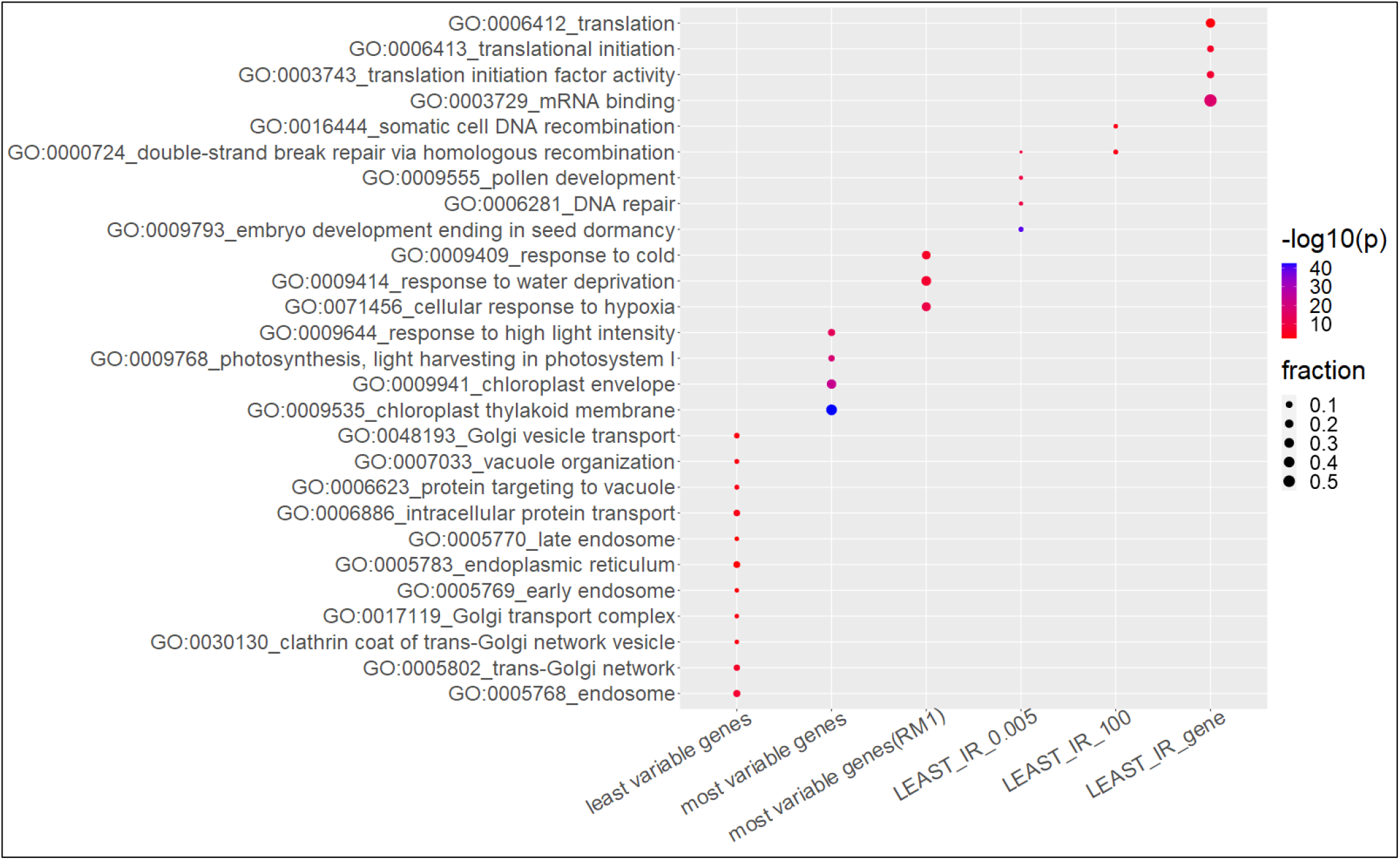
The GO enrichment plot of selected GO terms. Please refer to Suppl Tables 1∼9 for the complete GO enrichment reports of all gene lists. In this plot, points were colored according to -log_10_ p-values computed by using the TopGO elim method. Point sizes were corresponding to fractions of intersecting genes to sizes of input gene lists.

The combined findings on highly ranked genes in the stably expressed gene set are remarkably consistent across three GO categories, and they support the contention that reliable, steady expression of a cohesive set of genes involved in the above-mentioned cell biological processes are critical for normal physiological and developmental processes in 2-week-old seedlings.

#### 100 most variably (least stably) expressed genes (Table S3 and Table S4)

As might be expected from the known dependence of plant growth and development on light cues, GO enrichment analyses of variably expressed genes revealed a predominance of chloroplast and photosynthesis genes (**Figure 2** and **Table S3, GO-C, GO-F and GO-P**). When chloroplast and photosynthesis genes were removed from the GO analysis (**Table S4, RM1**), various stress-related genes, such as those involved in cellular responses to hypoxia, water deprivation and cold, were highly ranked (**Figure 2** and **Table S4, GO-P**).

These findings demonstrate that the expression of light and stress-responsive genes can readily fluctuate in two-week-old seedlings. This ability permits plants to adjust to rapid and unpredictable alterations in light intensity and environmental conditions.

### II. Identification and functional categorization of usually retained and usually spliced introns

We examined our RNA-seq data set for features of pre-mRNA splicing by identifying introns that were either usually retained or usually spliced, respectively, and functionally categorizing the respective genes by GO analyses.

#### Least stably retained (usually spliced) introns

##### Least_IR_0.005 (**Table S5**)

Dividing read depth of every intron by average depth of its neighboring exons, 4314 introns in 2105 genes had ratios less than or equal to 0.005 in all 50 RNA samples. This means that read depths of neighboring exons were at least 200 times higher than that of the respective introns in all of the 50 samples. GO_P analysis on this entire dataset revealed a possible significant enrichment of certain developmental genes in the category ‘embryo development resulting seed dormancy’ (rank 1) and ‘pollen development’ (rank 7). There were also some possibly interesting enrichments in DNA repair processes, such as ‘DNA repair’ (rank 4) and ‘double strand break repair via homologous recombination’ (rank 1) (**Figure 2** and **Table S5**). However, the importance of these results is not clear since members of these functional categories were not observed among highly ranked genes in GO_C and GO_F enrichments lists.

##### More refined analysis of least retained (usually spliced) introns (Least IR_100; **Table S6**)

For a more selective analysis, the 100 top ranked introns from the list of the stably retained/usually spliced introns in **Table S5** were used for GO analysis (**Table S6**). Although the developmentally related genes identified using the full dataset were not as pronounced in this limited dataset, a possible enrichment of genes involved in DNA repair again emerged: In the GO_P list, rank 3 is double strand break repair via homologous recombination and rank 4 is somatic cell DNA recombination (**Figure 2** and **Table S6)**. However, as for the more extensive analysis (previous paragraph), the importance of these results is not yet clear.

##### Least IR_gene **(Table S7)**

Thirty genes were identified as containing usually spliced introns. In other words, these genes were typically expressed without any intron retention in all 50 samples. There may be an interesting enrichment of translation-related genes in this group (**Figure 2** and **Table S7**, GO_P, GO_F), but apart from cytosolic ribosomal small subunit (GO:022627, rank 3) this trend was not convincingly observed in the GO_C class and hence is of uncertain significance.

#### Usually/mostly retained introns

##### *Most_IR_0.9 and Most_IR 100* **(Table S8 and Table S9)**

471 introns present in 438 genes showed retention ratios no less than 0.9 in all 50 samples. This indicates that more than 90% of the respective transcripts contain the entire intron in all of the 50 samples; i.e. more than 90% of the time, the intron was retained. Further ranking of these 471 introns were made based on their products of the 50 retention ratios. Notably, in both the 471-intron set (**Table S8**) as well as the top 100-intron set (**Table S9**), the first and last introns were the ones that fell into the usually-retained category. Of 471 introns that were usually retained (**Table S8**), 309 (65%) involved the first or last intron. Of these, 58 introns were the sole intron in single intron genes, and 56 introns belonged to double intron genes. If these single intron genes and double intron genes were excluded, 195 (54%) out of 357 (= 471-58-56) of the usually-retained introns were the first or the last introns. When considering only the top 100 usually-retained introns, 49 of them belonged to genes with three or more introns and 47 of these were the first and the last introns. In **Figure 3**, it was observed that usually retained introns from the MOST_IR_0.9 list (**Table S8**) showed a preferential location at the two ends of genes, whereas usually spliced introns from the LEAST_IR_0.005 list (**Table S5**) have a preferential location at the more internal central portions of genes.

**Figure 3:**
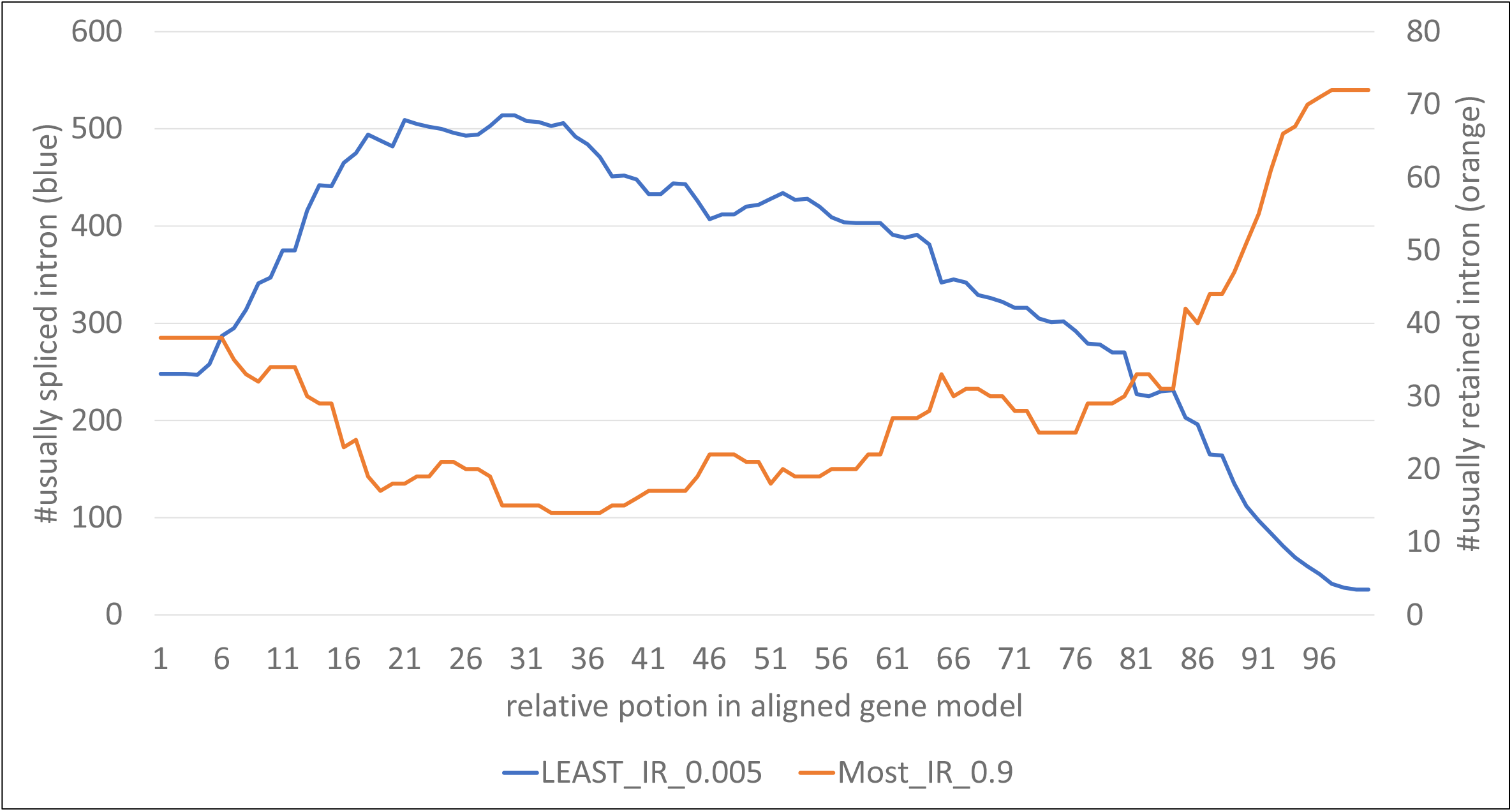
Positional trends of usually retained introns and usually spliced introns. Intron positions were all aligned to the same 100-slot array for all multi-intron genes. The curves are representing aggregated positional preferences in genes of given intron lists.

No consistent enrichments in any GO categories were observed when either the total dataset or the top 100 introns were subjected to GO enrichment analyses.

##### *MOST_IR_gene 0.7* **(Table S10)**

Only three genes containing introns with a retention ratio greater than 0.7 in all 50 samples are on the most retained IR list. Because of the very small sample size, a meaningful GO analysis was not possible.

### III. Features of the least variably (most stably), most variably (least stably) expressed genes, and genes containing usually spliced/retained introns (Figures 4 and 5)

To further characterize stably and variably expressed genes and introns, we analyzed whether there were any associations between stability/variability of expression and gene expression levels, gene length and intron number.

**Figure 4:**
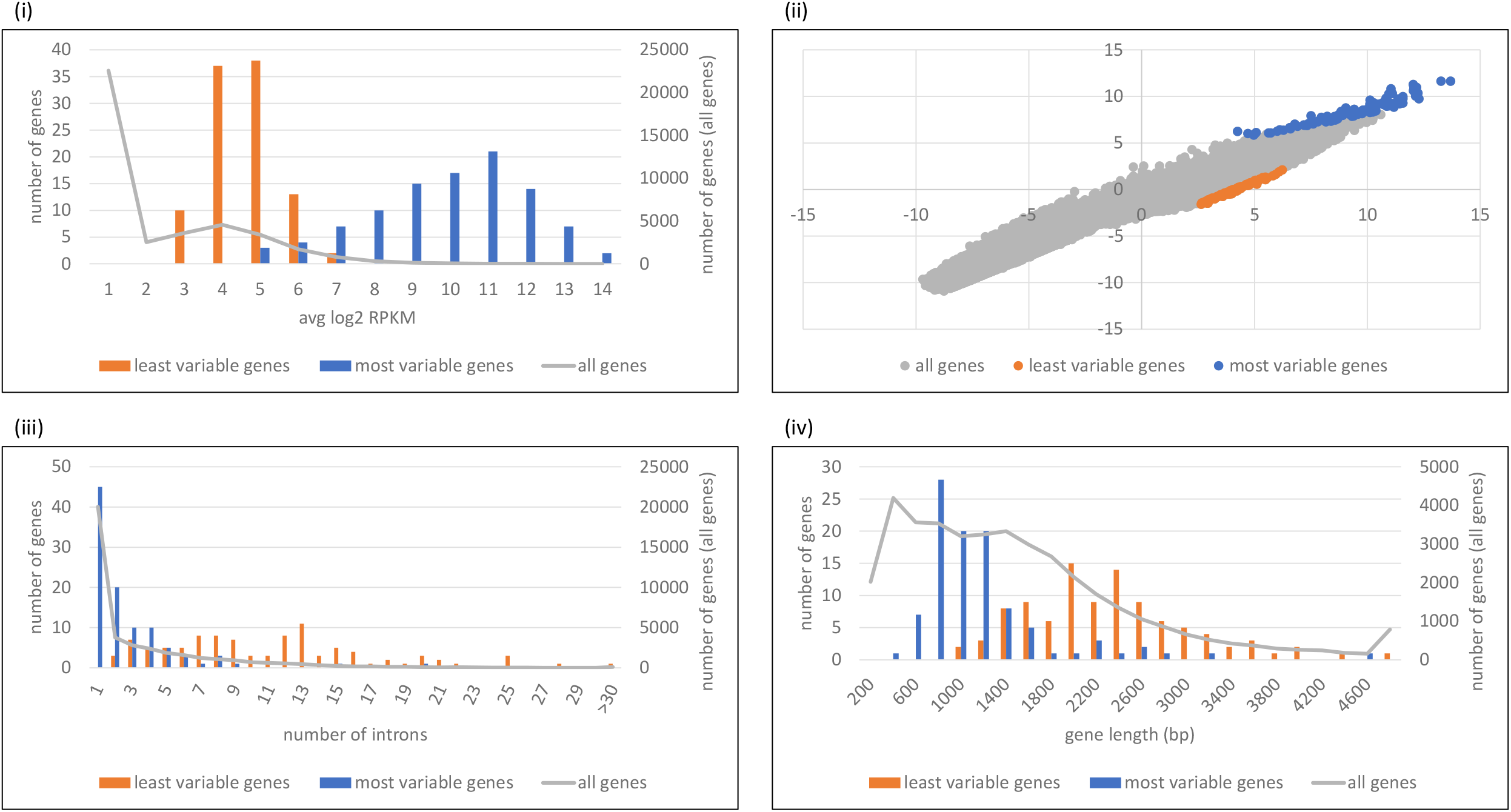
Distributions of least variable genes and most variable genes. (i) and (ii) Normalized expression levels and deviations of all genes and selected subsets. (i) Expression level distributions of least variable genes, most variable genes, and all genes. (ii) Expression level distribution of all genes in mean-stdev coordinates, in log2-log2 scale. Least variable genes and most variable genes were colored in orange and blue, respectively. (iii) and (iv) Distributions of intron numbers and gene lengths of least variable genes and most variable genes, respectively. The differences showed significance of p<2.2e-16 by two-sided KS test.

**Figure 5:**
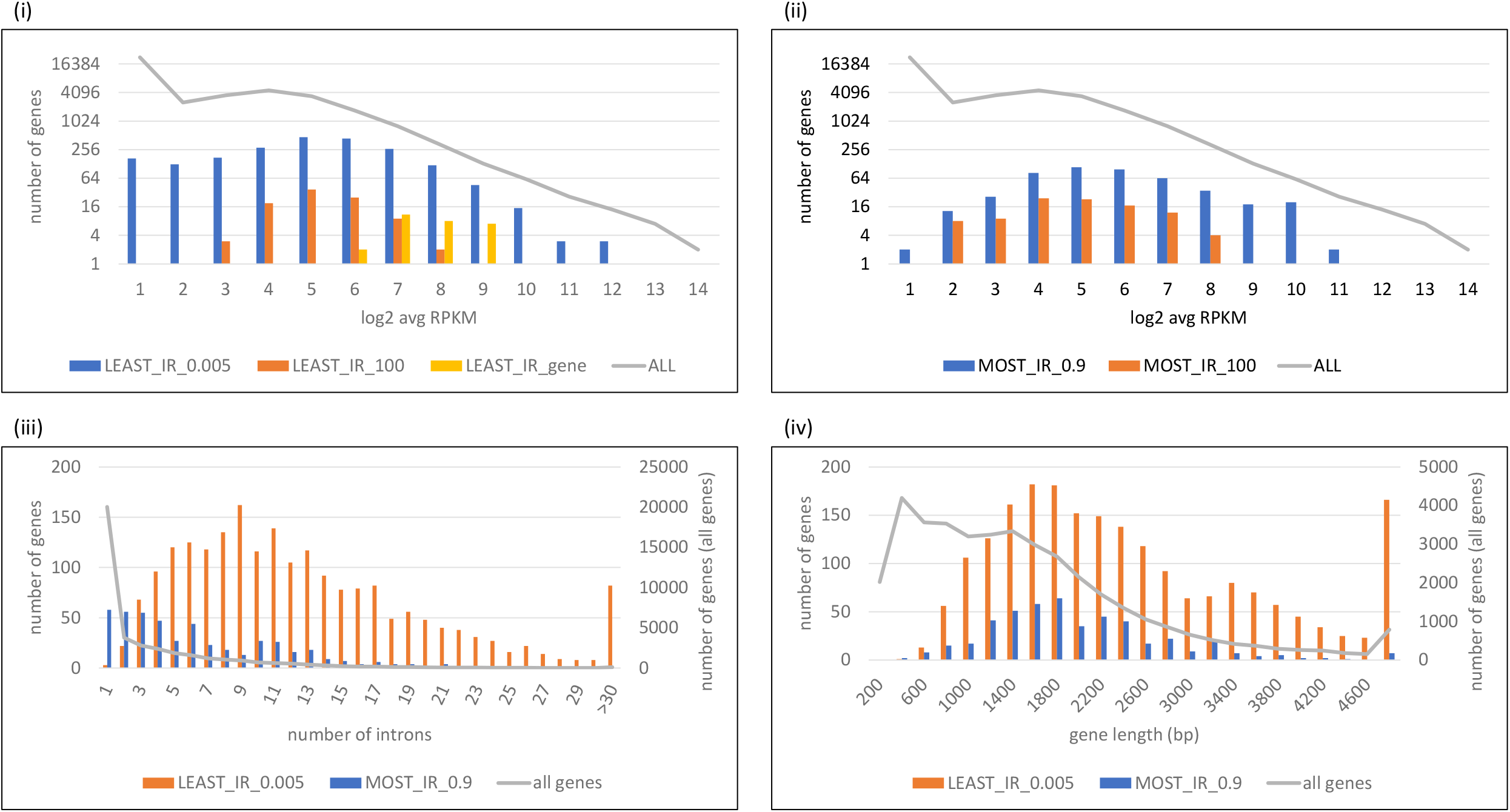
Expression level distributions of genes containing usually retained introns and usually spliced introns (i) Distributions of genes containing usually spliced introns. Note that the Y-axis is in log scale. Least IR gene sets: LEAST_IR_0.005 for genes with at least one intron with retention ratios less than 0.005 in all 50 samples, LEAST_IR_100 for genes of top 100 introns from LEAST_IR_0.005 with the 100 smallest products of retention ratios, and LEAST_IR_gene for multi-intron genes with >90% introns showing retention ratios less than 0.01 in all 50 samples. (ii) Distributions of genes containing usually retained introns. Note that the Y-axis is in log scale. Most IR gene sets: MOST_IR_0.9 for genes with at least one intron with retention ratios greater than 0.9 in all 50 samples, and MOST_IR_100 for genes of top 100 introns from MOST_IR_0.9 with the 100 greatest products of retention ratios. (iii) and (iv) Distributions of intron numbers and gene lengths of genes containing usually spliced introns and genes containing usually retained introns, respectively. The differences showed significance of p<2.2e-16 and p=4.463e-14 by two-sided KS test, respectively for intron number distributions and gene length distributions.

#### Gene expression level (avg log2 RPKM. Steady state RNA) vs. variability of expression (Figure 4i)

The expression level distributions in **Figure 4**i showed that the least variably (most stably) expressed genes are usually expressed at a median level whereas the most variably (least stably) expressed genes tend to be highly expressed. The mean-standard deviation plot (**Figure 4**ii, in log2-log2 scale) showed that the least variably and most variably expressed genes have lower and higher RPKM standard deviations, respectively, compared to genes of similar expression levels.

When observing the distributions of intron numbers and gene lengths for the least variably and the most variably expressed genes (**Figure 4iii** and **iv**), we noted that the least variably expressed genes contain more introns and are longer than the most variably expressed genes (p<2.2e-16, two-sided KS test). These results suggest that transcription and intron splicing are correlated with stabilized gene expression.

For genes containing usually spliced introns, we made the following observations (**Figure 5i**). First, for genes containing any intron that was usually spliced in the 50 samples, their expression level distribution is similar to that of the whole genome. For genes containing one of the top 100 usually spliced introns, the expression levels tended to be median. The expression levels of multi-intron genes whose introns were usually spliced ranged from median-to-high.

For genes with at least one usually retained intron (**Figure 5**ii), their expression level distribution tended to be median, as well as for genes with at least one of the top 100 retained introns. It was also noticed that the expression levels of some genes with at least one usually retained intron were slightly higher than those of genes with top 100 retained introns (p=0.029, one-sided KS test. **Figure 5**ii, blue bars in log2 RPKM 9∼11).

We also investigated the distributions of intron numbers and gene lengths of genes containing usually spliced introns (LEAST_IR_0.005) and those containing usually retained introns (MOST_IR_0.9) (**Figure 5iii** and **iv**). Notably, genes in LEAST_IR_0.005 contains significantly more introns than MOST_IR_0.9 (p<2.2e-16, two-sided KS test), whereas genes in LEAST_IR_0.005 are longer than genes in MOST_IR_0.9 at a less significant level (p= p=4.463e-14, two-sided KS test). These results suggest that genes containing usually spliced introns or usually retained introns is correlated with transcript lengths and intron numbers.

#### Summary

Low variability (high stability) of expression was often observed for genes with median expression levels, longer gene lengths and higher numbers of introns compared to the most variably (least stably) expressed genes. The biological significance of these results and what they indicate in terms of possible mechanistic aspects of the control of gene expression are not yet clear but can be explored in future investigations.

## Discussion

The aim of our study was to identify and functionally categorize the most stably and the most variably (least stably) expressed genes and retained introns in young Arabidopsis seedlings by analyzing a large, uniform dataset comprising 50 transcriptomes. To our knowledge, a similar large-scale study on stability of gene expression and intron retention has not been performed previously in plants.

Two notable findings emerged from our study. First, unlike the variable expressors, which perhaps predictably identified known or presumed environmentally-sensitive genes, it was not clear a priori what to expect from the analysis of the most stably expressed genes. One might anticipate that this group includes genes having an essential role in growth and development of two-week old Arabidopsis seedlings, such as certain cell-type-specific transcription factors or ion channel proteins that may be particularly required at this stage. Structural genes encoding cytoskeletal proteins, e.g. actin and tubulin, and other housekeeping genes (e.g. ribosomal proteins), which are needed in most cell types, may also be among the stable expressors in these seedlings. Indeed, some members of these protein families are often used as an internal standard for RNA quantification by PCR (Yu *et al*., 2019).

However, these general classes were not among the most highly ranked GO categories in the stably expressed gene set that we identified. Rather, genes encoding endocytic and secretory pathway proteins as well as proteins of the Golgi complex and endoplasmic reticulum were the most strongly represented among the highest ranked categories of stable expressors. Endosomes are a diverse set of intracellular sorting organelles that regulate trafficking and exchange of proteins and lipids among cellular compartments of the endocytic and secretory pathway, in particular the plasma membrane, endoplasmic reticulum, Golgi, trans Golgi network and vacuoles/lysosomes (Otegui *et al*., 2010). Endocytosis permits removal of proteins from the cell membranes or uptake of soluble material from the external environment. The secretory pathway conveys proteins and lipids to the cell membrane and to intracellular organelles. The cellular machineries and mechanisms that govern molecular trafficking along the endocytic pathway are evolutionarily conserved and hence a fundamental and defining feature of eukaryotic cells (Schmid *et al*., 2014).

The second notable finding from our study concerns the locations of highly retained or usually spliced introns within a pre-mRNA. Untranslated regions (UTRs) at the 5’ and 3’ ends of a pre-mRNA are essential for modulating transcription and translation. Although not often considered in terms of the presence of introns (Bicknell *et al*., 2012), UTRs in these locations can have significant functional consequences for gene regulation. The presence of an intron in 3’ UTRs can regulate transcript stability whereas introns in 5’UTRs can influence promoter selection, the mechanism of mRNA transport and cellular localization of mature mRNAs (Bicknell *et al*., 2012). Introns in UTRs thus represent regulatory modules that are potentially removable by splicing.

Two caveats of our study are important to mention. First, our experiments examined steady state RNA transcript levels, which are the result of both transcription rates and RNA turnover. The extent to which these two processes either separately or in combination influenced our findings is not yet known. Second, the two-week old Arabidopsis seedlings used in our study do not comprise a single cell type but are a mixture of different cells types. A significant proportion of these cells constitute meristematic regions at the shoot and root apices. Meristems contain undifferentiated stem cells that continuously divide to replenish the stem cell populations and to produce new cells that develop into the aerial and underground parts of the growing plant. Hence, they are sites of significant developmental activity

In summary, our results demonstrate the power of using large data sets to uncover previously underappreciated patterns and trends of gene expression. The stably expressed genes we identified make up a functionally cohesive set of genes that have essential roles – via the endocytic and secretory pathways - for the structural and functional organization of the cell. Clearly the steady and reliable expression of genes acting in these pathways are crucial for plant development and viability. In addition to their inherent interest for cell biological phenomena, the identification of regulatory elements of stable expressors such as transcriptional promoters and UTRs that may be regulatable by intron splicing, can potentially be useful for applications in plant gene technology that require reliable expression of introduced genes (Zhou *et al*., 2016; Yang *et al* (2023).

## Data Displays

**Figure 1**: Sample sizes

**Figure 2**: Selected GO categories

**Figure 3:** IR position

**Figure 4:** Least and most variable genes

**Figure 5:** Least and most Irs

## Supplementary Tables

**Table S1**: Read mapping statistics

**Table S2**: Gene least variable (most stable)

**Table S3**: Gene most variable

**Table S4**: Gene most variable RM1

**Table S5**: Least IR 0.005

**Table S6**: Least IR 100

**Table S7**: Least IR gene

**Table S8**: MOST IR 0.9

**Table S9**: MOST IR 100

**Table S10**: MOST IR gene 0.7

